# Cortical tracking of postural sways during standing balance

**DOI:** 10.1101/2023.12.08.570752

**Authors:** Thomas Legrand, Scott J. Mongold, Laure Muller, Gilles Naeije, Marc Vander Ghinst, Mathieu Bourguignon

## Abstract

Maintaining an upright stance requires the integration of sensory inputs from the visual, vestibular and somatosensory-proprioceptive systems by the central nervous system to develop a corrective postural strategy. However, it is unclear whether and how the cerebral cortex monitors and controls postural sways. Here, we asked whether postural sways are encoded in ongoing cortical oscillations, giving rise to a form of corticokinematic coherence (CKC) in the context of standing balance. Center-of-pressure (CoP) fluctuations and electroencephalographic cortical activity were recorded as young healthy participants performed balance tasks during which sensory information was manipulated, by either removal or alteration. We found that postural sways are represented in ongoing cortical activity during challenging balance conditions, in the form of CKC at 1–6 Hz. Time delays between cortical activity and CoP features indicated that both afferent and efferent pathways contribute to CKC, wherein the brain would monitor the CoP velocity and control its position. Importantly, CKC was behaviorally relevant, as it predicted the increase in instability brought by alteration of sensory information. Our results suggest that human sensorimotor cortical areas take part in the closed-loop control of standing balance in challenging conditions. Importantly, CKC could serve as a neurophysiological marker of cortical involvement in maintaining balance.

## Introduction

Evolution has led humans to adopt an upright posture, thanks to—and fostering—the development of an exceptionally large brain as compared to other species. As a corollary, the cortical and subcortical network supporting motor control is complex, and even the simplest tasks, such as maintaining upright balance, require the involvement of the spinal cord and brainstem for reflexive control, and the cerebral cortex for adjustments to the environment.^1–5^ In this regard, sensory inputs from the visual, vestibular and somatosensory-proprioceptive systems to the cortex play a key role in the control of standing balance.^6^ The integration of this sensory information by the cortex is at the basis of the development of a postural corrective strategy. Our current understanding of the cortical mechanisms implicated in balance is based on the analysis of surrogate variables, such as body sway measurements, while the proprioceptive, visual or vestibular systems are manipulated. This led to the description of the phenomenon known as sensory weighting and reweighting,^7,8^ where the contribution of each sense to balance depends on and evolves with its reliability. However, the direct study of the integration of these sensory cues by the cortex during balance tasks has proven challenging.

From the biomechanics perspective, balance maintenance is achieved by adjusting the position of the center of pressure (CoP) on the support surface.^9^ This means that when the center of mass is displaced from its neutral position, changes in CoP are made to redirect it back toward the neutral axis. Indeed, previous studies have shown that the CoP regularly oscillates along the antero-posterior axis about the vertical projection of the center of mass on the ground during quiet standing.^10,11^ Whether and how the cerebral cortex monitors and controls CoP fluctuations is still unsettled.

Studies using electroencephalography (EEG) have identified changes in cortical oscillatory dynamics across several frequency bands during quiet stance compared with sitting. ^12–14^ Hypothetically, these changes at the cortical level would reflect the engagement of mechanisms responsible for modulating muscle contractions over time, with the aim to regulate balance. Further supporting this view, the cortical oscillatory dynamics was modulated by the quality of available visual and proprioceptive cues.^3,14,15^ Indeed, the manipulation of the sensory information during a balance task, by either removal or alteration, makes balance control significantly more challenging due to increased difficulty in estimating the dynamics of the body, leading to an increase in body sway,^16,17^ and cortical involvement.^18^ Besides, cortico-muscular coherence (CMC) has previously been used to assess the cortical processes for postural maintenance.^2,4^ However, the nature of this coupling is still unclear,^19,20^ and it does not inform about how postural sways are integrated in the central nervous system.

A study using electroencephalography (EEG) has identified cortical activity time-locked to naturally occurring postural sway in the form of an evoked negative potential ∼100 ms prior to the onset of postural reactions during quiet standing with eyes closed ^3,13^. In view of its timing, this cortical activity seemed to reflect the engagement of efferent mechanisms responsible for modulating muscle contractions over time, with the aim to regulate balance ^3,13^. This discrete evoked potentials seen in relation to a particular feature of the postural sway could underscore the existence of a more continuous monitoring process of postural sway. A promising means to gain insight into this process is corticokinematic coherence (CKC). CKC, the coupling between brain activity and movement kinematics,^21,22^ has been shown to specifically reflect the integration of proprioceptive input by the cortex during active and passive finger movements.^20,23–25^ Thus far it has been studied only for repetitive movements of the upper or lower extremity in a sitting position. Therefore, we expect CKC to bring further insights into the sensory integration processes taking place during balance.

This study investigates postural maintenance through the lens of CKC, to determine if postural sways are encoded in ongoing cortical oscillations. Given that cortical involvement is expected to depend on the complexity of the balance task, we hypothesized that the extent of cortical sway encoding increases with the suppression or alteration of visual and proprioceptive input. In addition, we aimed to determine which aspects of the sways are encoded, focusing on excursion and velocity of the CoP. We then tested the hypothesis that cortical sway encoding hinges on both afferent and efferent signaling, in line with the dominant afferent contribution commonly reported during upper limb voluntary movements and the efferent nature of the evoked potential recorded prior to a postural reaction. And finally, we asked whether such sway-driven CKC is relevant for postural stability.

## Results

To study CKC during postural maintenance, a group of 36 young healthy adults maintained bipodal quiet standing on a solid surface or on a foam mat with eyes open or closed, leading to 4 conditions (*solid-eyes-open, foam-eyes-open, solid-eyes-closed*, and *foam-eyes-closed*), each performed for a total of 10 minutes (Figure 1A-C). Cortical activity was recorded with EEG, along with the position of the CoP with a force plate. CoP traces were filtered through 0.1-10 Hz.

**Figure 1.**
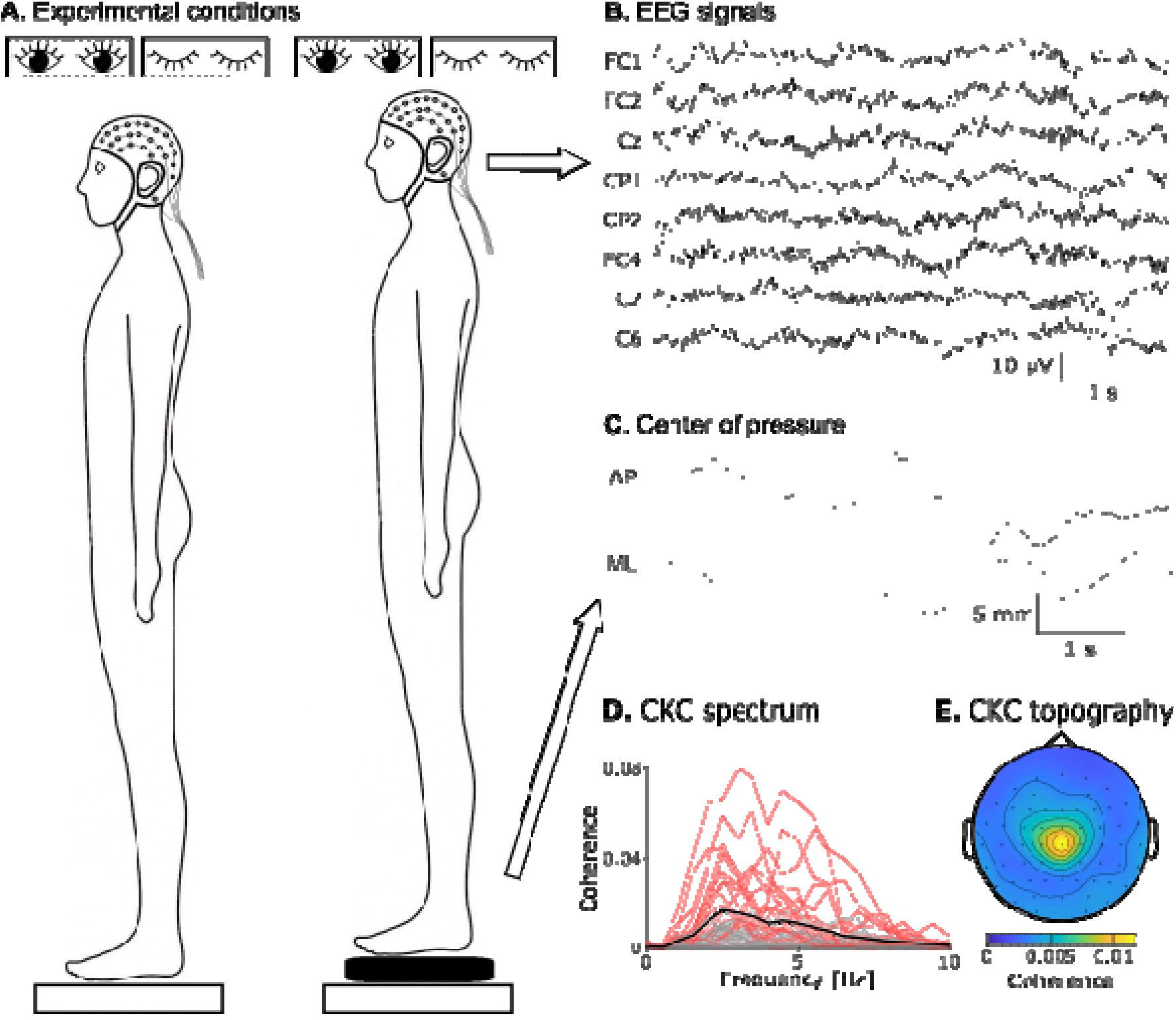
Experimental set-up and CKC estimation. A — Participants were equipped with a 63 channel EEG-cap and stood on a force plate in 4 experimental conditions: either on a hard surface or on foam pads, and with eyes open or closed. B — Six second excerpt of EEG collected during the foam-eyes-closed condition. C — CoP signals along the mediolateral (ML) and antero-posterior (AP) axes. D — CKC spectrum at the electrodes overlying the lower limb primary sensorimotor cortex for CoP velocity when standing on foam eyes closed. There is one trace for each participant with (red traces) and without (gray traces) significant CKC and one for their group average (black traces). For each participant, the CKC spectrum was taken from the best electrode from among a predefined subset of electrodes overlying the lower limb primary sensorimotor cortex. E — Topography of the CKC for CoP velocity in the band 1-6 Hz, averaged across the entire cohort and conditions.

### CoP variability

The inspection of CoP traces obtained in the four different balance conditions indicated that medio-lateral (ML) and antero-posterior (AP) sways were larger when standing on a foam surface compared with a solid surface, and when keeping the eyes closed compared to open (see Figure 2A). To objectivate these observations, we estimated three instability parameters consisting of the mean speed and standard deviation (SD) of the position in the ML and AP direction (see Figure 2B) and an ANOVA identified a significant effect of the balance condition on these parameters (speed, *F*_*3,105*_ = 49.7, *p* < 0.0001; ML SD, *F*_*3,105*_ = 284, *p* < 0.0001; AP SD, *F*_*3,105*_ = 210, *p* < 0.0001). Overall, pairwise comparisons between conditions for the SD of ML and AP position fluctuations confirmed our initial observation of increased instability when standing on a soft surface and when closing the eyes with only a few exceptions (see Figure 2B for the result of post-hoc comparisons), in line with previous studies. As for speed, it was significantly increased in the most challenging condition compared with the three other conditions within which no significant differences were observed. The absence of significant difference in average speed among these later 3 easiest conditions could be due to the imposed position of the feet at 20 degrees, providing further stability than a strictly narrow stance. The contrast between average speed and the variability of position could be explained by a difference in sensitivity to the balance condition for these variables.

**Figure 2.**
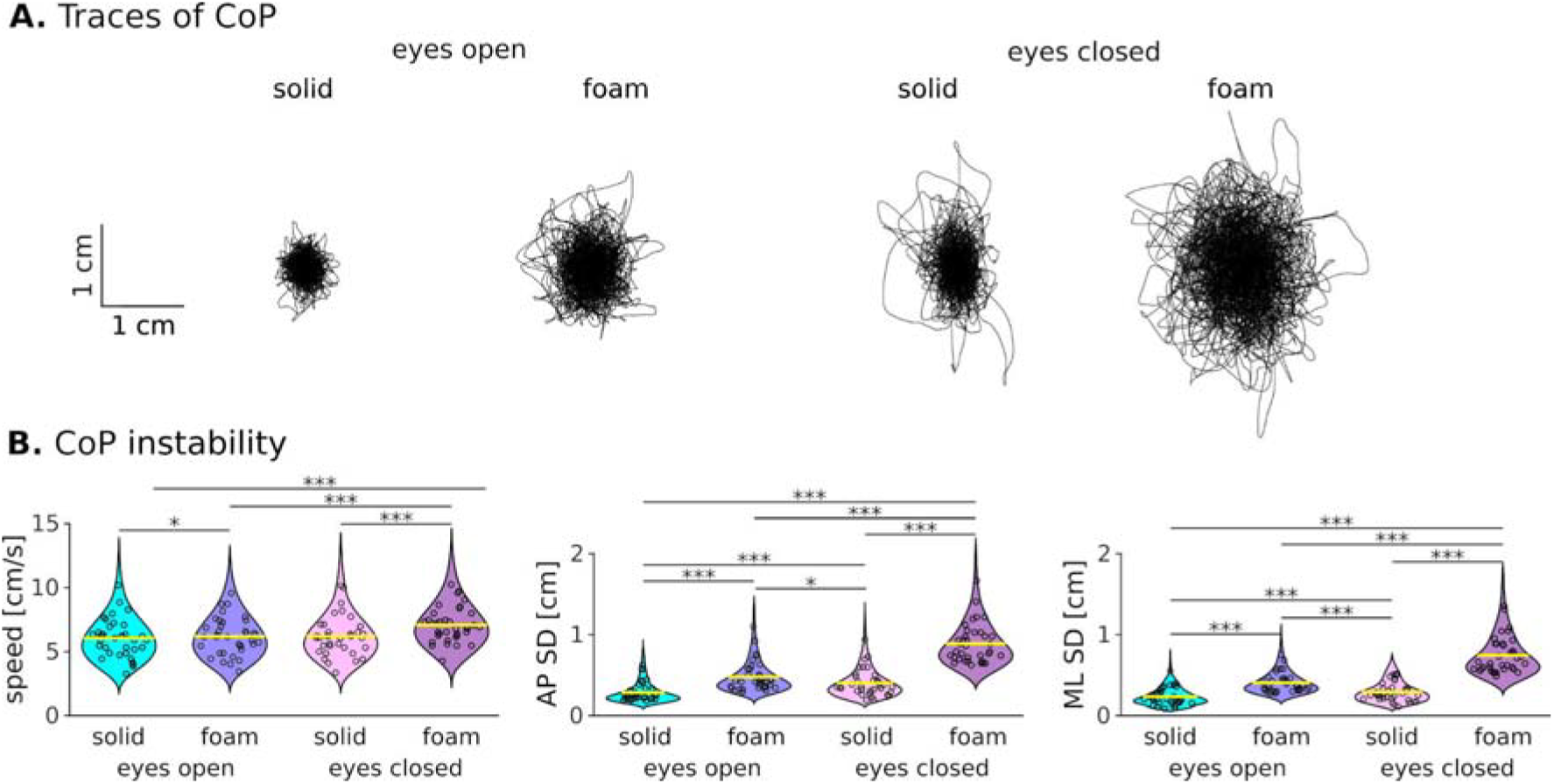
Effect of conditions on postural sways. **A** — Traces of the center of pressure (CoP) of a typical participant during a 5-min trial for each balance condition. **B** — Distribution of the instability parameters (CoP speed, medio-lateral (ML) standard deviation (SD) and antero-posterior (AP) SD) for each balance condition. Each individual subject’s value is indicated with circles. Significant differences between conditions are indicated with stars (*, p < 0.05; **, p < 0.01; ***, p < 0.001).

### Neural encoding of CoP

Multiple CoP features were assessed for their coupling with EEG signals. These features were the velocity along the ML (vCoP_ML_) and AP directions (vCoP_AP_), and the vector sum of position (rCoP) and velocity (vCoP). Importantly, these CoP features appeared relatively decoupled from one another. Indeed, coherence between feature pairs averaged across participants were all below 0.05 in the 1–6 Hz range, except for the pair rCoP–vCoP where coherence was above 0.2 at frequencies below 1.5 Hz and reached about 0.1 in between 2–6 Hz (see Figure 3). Therefore, CKC was assessed between each EEG signal and each of these CoP features. For further analyses, we retained only the electrode from among those typically overlying the lower limb primary sensorimotor cortex (SM1) (Cz, C1, C2, FCz, FC1, FC2, CP1, CP2) that yielded the highest CKC averaged across 1–6 Hz and balance conditions. From this point on, it is referred to as EEG_SM1_.

**Figure 3.**
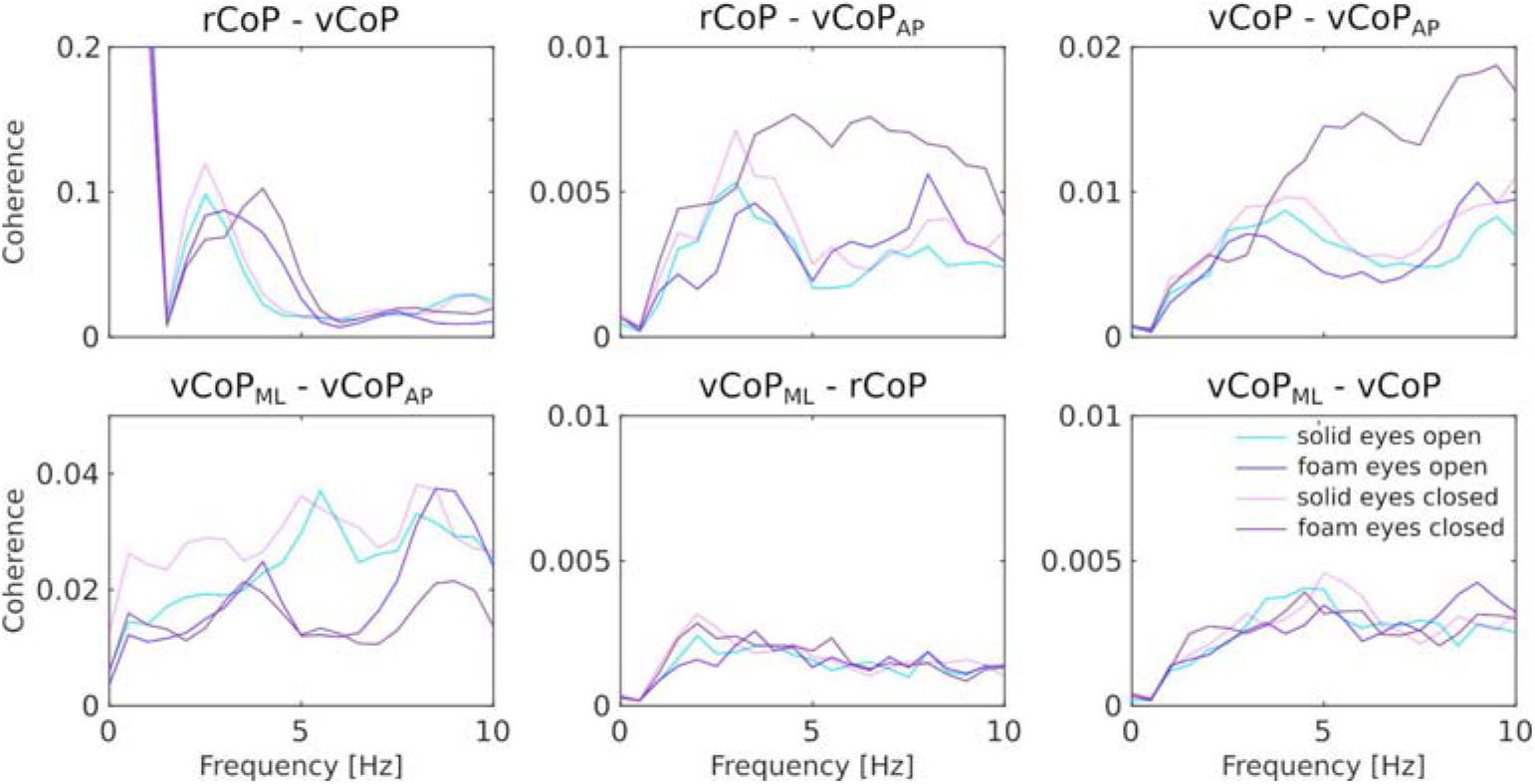
Coherence between CoP features. Displayed is the mean across all participants for each CoP feature pair and each condition.

First of all, the balance condition did not affect the number of artifact-free epochs used in the analyses (F_3,105_ = 0.88, *p* = 0.45; mean ± SD epochs across conditions and participants, 1100 ± 20). This ensures CKC can be validly compared between conditions.

Table 1 presents the number and proportion of participants showing significant CKC for each condition and feature type. Overall, this proportion was significantly higher than expected by chance in all conditions and for all CoP features, but it increased with the difficulty of the balance condition. The highest proportion was reached for vCoP in the most challenging condition (*foam-eyes-closed*). We therefore concentrate first on the results for this specific feature, then assess the effect of condition on CKC for all CoP features.

**Table 1.**
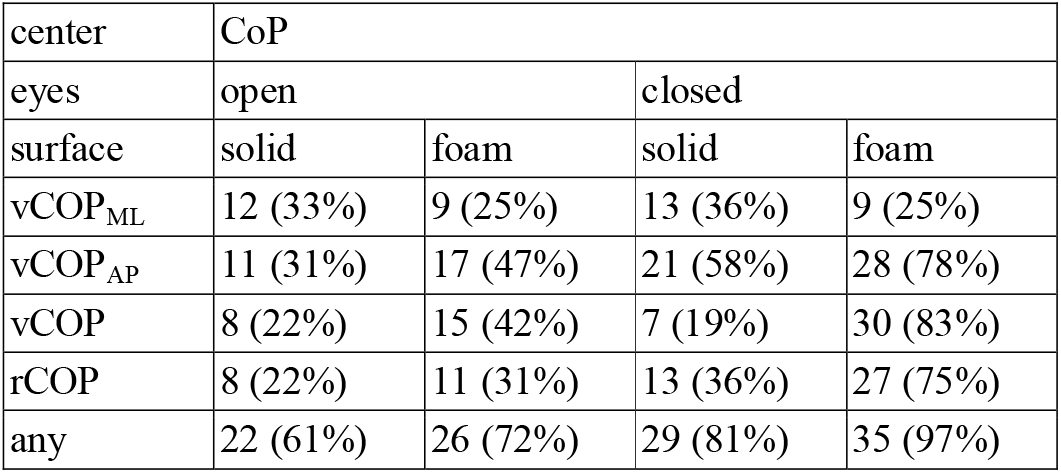
Number and percentage of participants with significant CKC with each feature, and with any of them.

| center | CoP |  |  |  |
| --- | --- | --- | --- | --- |
| eyes | open |  | closed |  |
| surface | solid | foam | solid | foam |
| vCOP<sub>ML</sub> | 12 (33%) | 9 (25%) | 13 (36%) | 9 (25%) |
| vCOP<sub>AP</sub> | 11 (31%) | 17 (47%) | 21 (58%) | 28 (78%) |
| vCOP | 8 (22%) | 15 (42%) | 7 (19%) | 30 (83%) |
| rCOP | 8 (22%) | 11 (31%) | 13 (36%) | 27 (75%) |
| any | 22 (61%) | 26 (72%) | 29 (81%) | 35 (97%) |

Figure 1D-E presents the spectra and topography of CKC for vCoP on foam eyes closed. Therein, CKC tended to peak in between 1 and 6 Hz. At the group level, the electrode showing the maximum level of CKC was Cz. This electrode was also the one where CKC was maximum in 81% of the participants with significant CKC.

Importantly, significant CKC in the most challenging condition (*foam-eyes-closed*) was uncovered in more than 75% of the participants for 3 of the CoP features (rCoP, vCoP_AP_ and vCoP). In all instances, group-averaged CKC magnitude tended to peak around Cz, and was well within our sensor selection covering SM1 (see Figure 4).

**Figure 4.**
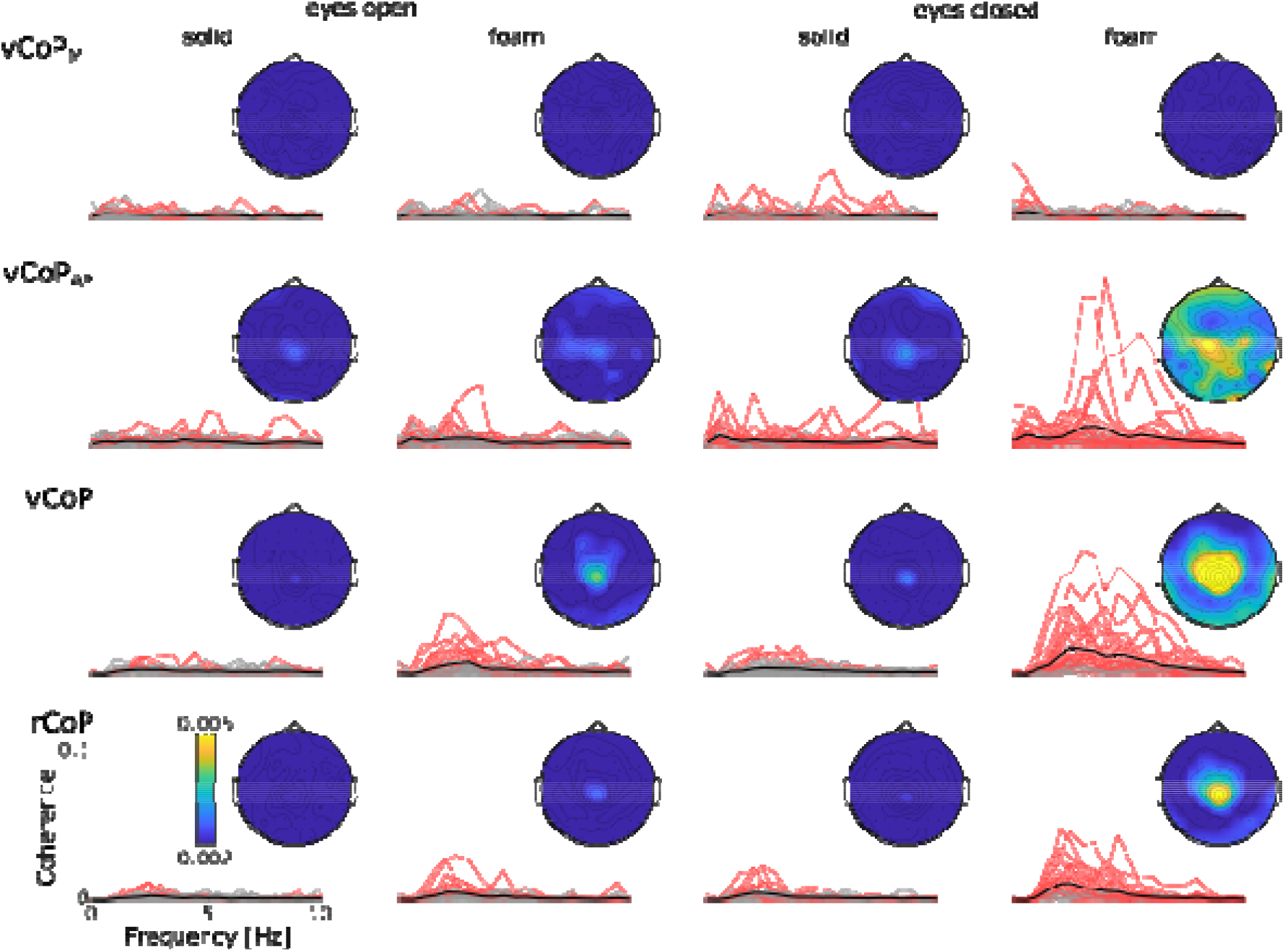
CKC spectrum for each CoP feature in each balance condition at the EEG_SM1_ electrode for each participant with (red traces) and without (gray traces) significant CKC and their group average (black traces). On the top right corner of each graph is the scalp distribution of the CKC values averaged across participants and across 1–6 Hz.

Figure 5 presents the distribution of individual CKC values (averaged across 1–6 Hz) for the 4 CoP features in all conditions. Visually, the pattern of variation across conditions of CKC for the 3 features highlighted in the previous paragraph (vCoP, vCoP_AP_ and rCoP) followed that of CoP instability, being higher when standing on a foam surface compared with a hard surface, and when keeping the eyes closed compared to open. In support of that observation, one-way ANOVAs run for each CoP feature separately identified a significant effect of condition on CKC for these 3 features (vCoP, *F*_3,105_ = 30.7, *p* < 0.0001; vCoP_AP_, *F*_3,105_ = 10.3, *p* < 0.0001; rCoP, *F*_3,105_ = 29.9, *p* < 0.0001), and not for the remaining one (vCoP_ML_, *F*_3,105_ = 0.22, *p* = 0.30). However, unlike CoP instability, the effect of the foam surface or closing the eyes in isolation appeared rather small, and only their combination gave rise to a marked increase in CKC (see also Figure 5 for post-hoc comparisons).

**Figure 5.**
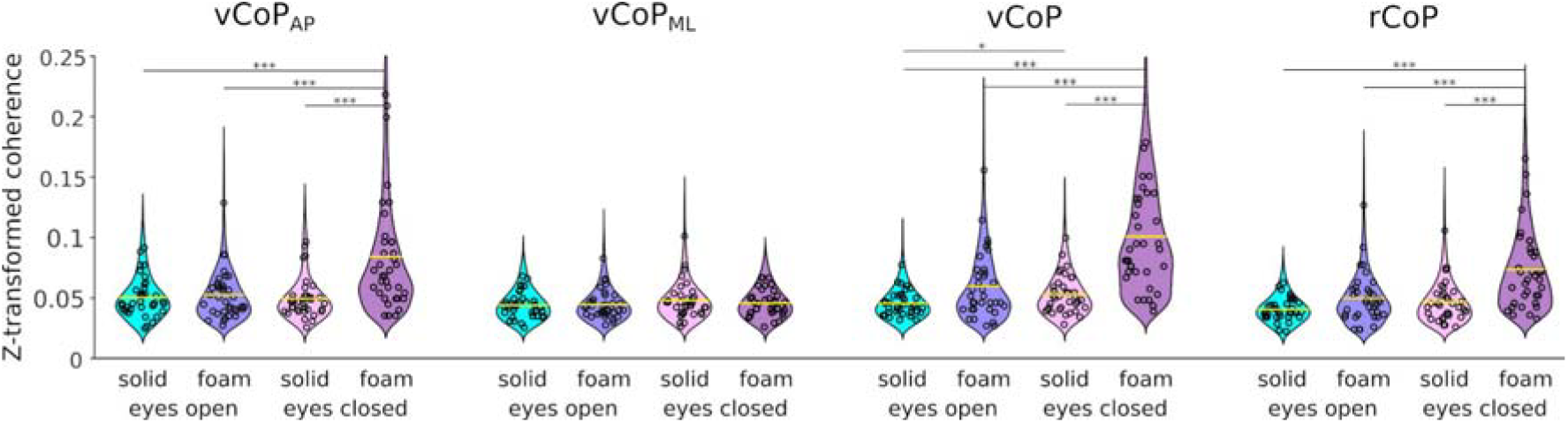
Distribution of CKC values for each CoP feature in the four balance conditions. Displayed are the CKC values averaged across 1–6 Hz and z-transformed (by means of the arctangent of the square-root). Each individual subject’s value is indicated with circles. Significant differences between conditions are indicated with stars (*, p < 0.05; **, p < 0.01; ***, p < 0.001).

We further estimated the peak amplitude of the EEG_SM1_ signal underlying the observed significant CKC values. The average amplitude across feature types ranged in 0.45– 1.09 μV (rCoP, 0.91 ± 0.32 μV; vCoP_ML_, 0.45 ± 0.13 μV; vCoP_AP_, 0.82 ± 0.29; vCoP, 1.09 ± 0.48 μV), in line with the amplitude of ∼1 μV previously reported for sway-based evoked responses.^3,13^

### Delays

The time delays between EEG_SM1_ and CoP features were estimated from the phase slope of their cross-spectral density, using only the data of participants with significant coherence, at participant-specific frequencies where significant coherence was found. A negative-phase slope corresponds to a positive delay whereby CoP variation precedes EEG_SM1_ activity. We performed these analyses only for CoP features yielding significant CKC in at least 20 participants in the most challenging condition (*foam-eyes-closed*). This led us to consider only rCoP, vCoP_AP_ and vCoP.

Figure 6 presents the evolution of the phase-frequency plots at the basis of the estimation of the phase delay between CoP features and EEG_SM1_. These phase delays were significantly different from 0 for rCoP (*t*_26_ = -5.22, *p* < 0.001) and for vCoP_AP_ (*t*_27_ = 2.17, *p* = 0.039) and were not significantly different from 0 for vCoP (*t*_29_ = 0.15, *p* = 0.88). The delays for rCoP were negative, with an average of -47 ± 47 ms (mean ± SD), indicating that EEG_SM1_ activity led fluctuations in rCoP. The delays for vCoP_AP_ were positive, with an average of 44 ± 108 ms (mean ± SD), indicating that fluctuations in vCoP_AP_ led EEG_SM1_ activity.

**Figure 6.**
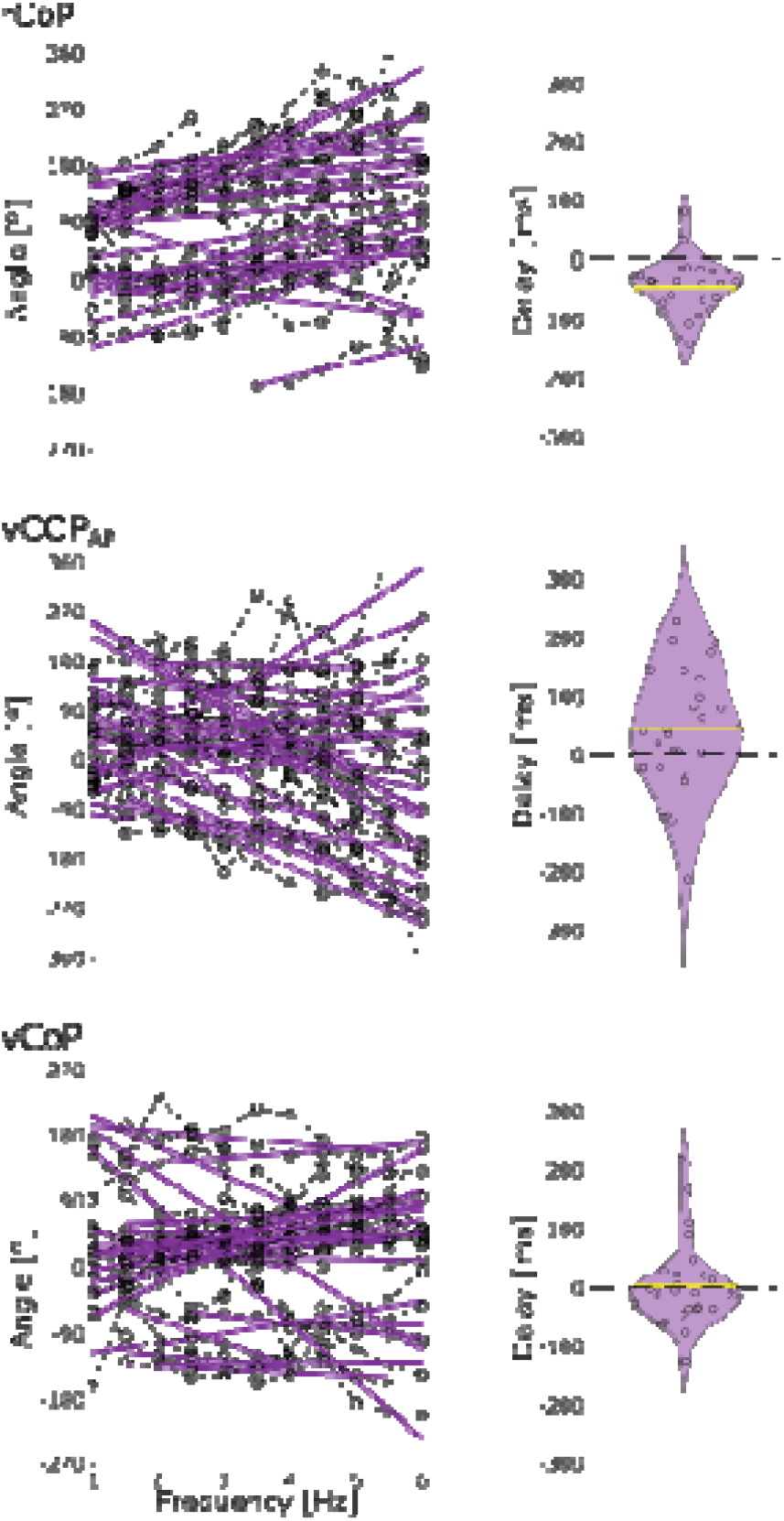
Estimated phase delays underlying the cortical encoding of postural sways when standing on foam eyes closed. **Left** — Phase frequency plots for the cross-spectrum between CoP features (rCoP, vCoP_AP_ and vCoP) and EEG_SM1_. Only data points of significant CKC are indicated, and connected within participants. A regression line per participant displaying significant CKC is shown in purple. **Right —** Violin plot of the delays derived from the slope of the regression lines. Positive delays indicate that fluctuations in the CoP feature precede those of cortical activity. Individual participant values are displayed with a circle.

To ascertain that differences in delays for rCoP and vCoP_AP_ reflect genuine differences in the nature of the cortical encoding of these features as opposed to a mere delay in between them, we also analyzed the delay between rCoP and vCoP_AP_ (see Figure 7). The average estimated delay was -17 ± 112 ms and was not found significantly different from 0 (*t*_26_ = -0.90; *p* = 0.37).

**Figure 7.**
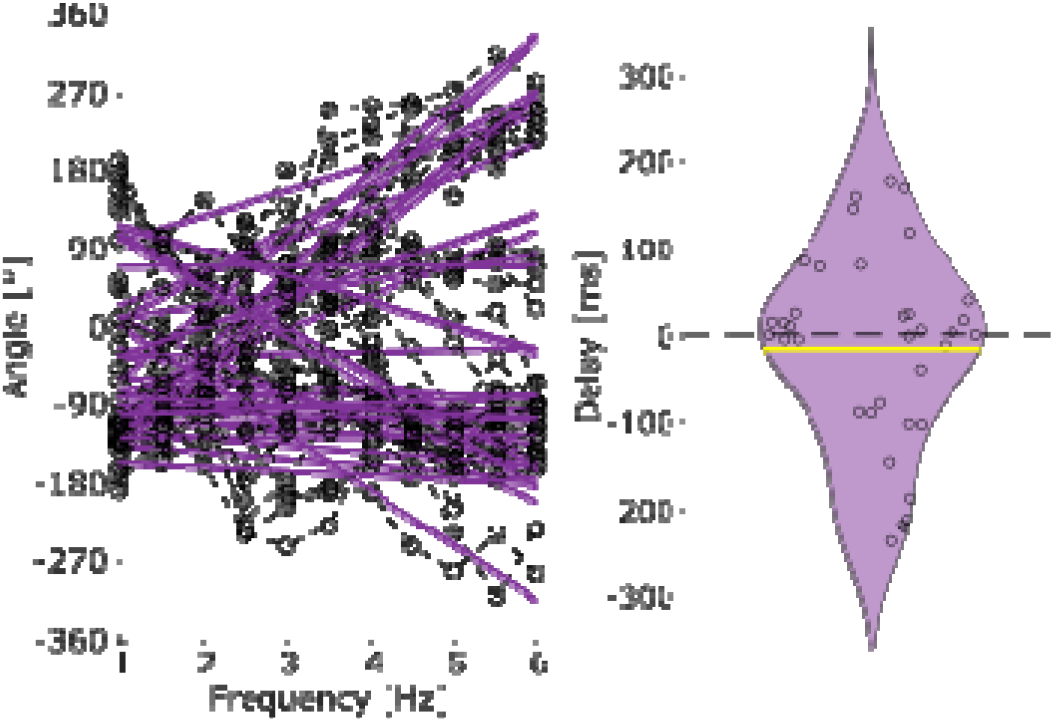
Estimated phase delays between rCoP and vCoP_AP_ when standing on foam eyes closed. **Left** — Phase frequency plots for the cross-spectrum between rCoP and vCoP_AP_. A regression line per participant is shown in purple. **Right** - Distribution of the delay between rCoP and vCoP_AP_ in the range 1-6 Hz when standing on foam with the eyes closed. Positive delays indicate that fluctuations in vCoP_AP_ precede those in rCoP. Individual participant values are displayed with a circle.

### Behavioral relevance of CoP cortical encoding

Our results identified significant CKC, especially for rCoP, vCoP and vCoP_AP_ in the most difficult condition. We therefore sought to determine the relevance of these couplings for balance maintenance, quantified with the SD of CoP_AP_ and CoP speed. Given the multivariate nature of the brain and balance data, we used regularized canonical correlation analysis (CCA). This analysis did not reveal a significant association between instability parameters and CKC values in the most challenging condition (*p* = 0.77). Next, we assessed whether CKC could instead be associated with the degree of increase in instability induced by standing on foam eyes closed, with the rationale that such degree reflects that to which balance is compromised compared with an individual’s typical reference point, begging for commitment of central resources to prevent falls.

Figure 8 presents the results of a regularized CCA applied to CKC and CoP variables, after each CoP variable in the most challenging condition were normalized by the value in the least challenging condition as follows:

**Figure 8.**
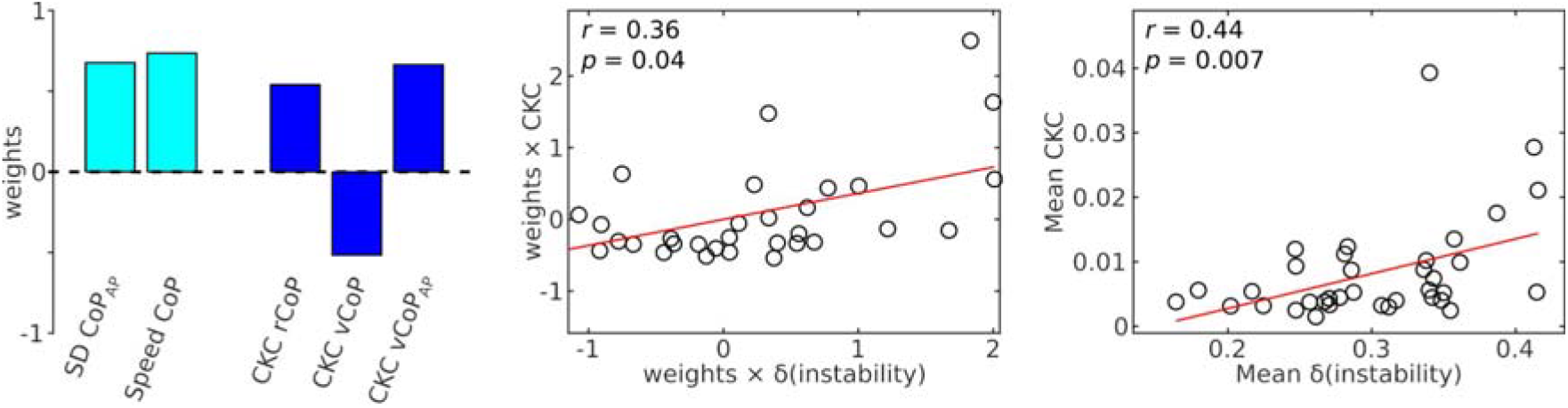
Behavioral relevance of the cortical encoding of postural sways as assessed with a CCA between CKC values for the 3 dominant features and the relative increase in CoP instability parameters. The relative increases, indicated with a δ, represent a contrast between the foam-eyes-closed and solid-eyes-open conditions. From left to right are the weights for the instability and CKC variables, the scatter plot of CKC variables combined according to these weights as a function of the instability parameters combined likewise, and the scatter plot for the mean of CKC across the 3 features as a function of the mean of the two instability parameters.

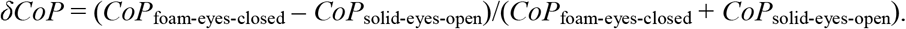

Such contrast was used because it is specific to changes induced by a reduction of sensory cues and factors out the inter-individual variability in the level of instability. It takes values between –1 and 1, with positive values indicating an increase due to reduction of sensory cues. It was not applied to CKC variables since these were mostly non-significant in the least challenging condition. This analysis identified a significant association (*p* = 0.04), characterized by CCA weights that were all positive and of similar amplitude, except for CKC for vCoP. This indicates a positive association between overall increase in instability and a contrast involving positive weights for CKC for rCoP and vCoP_AP_ and a negative weight for CKC for vCoP. However, CKC for vCoP was moderately and positively correlated with CKC for rCoP (*r* = 0.55, *p* = 0.0006) and CKC for vCoP_AP_ (*r* = 0.55, *p* = 0.0007), the latter two showing an even stronger correlation (*r* = 0.78, *p* < 0.0001). In addition, the contrast for CKC identified by CCA largely correlated positively with a more intuitive average across the 3 features (*r* = 0.84, *p* < 0.0001). As a result, the relative increase in instability averaged across the 2 instability features was significantly correlated with the CKC averaged across the 3 retained CoP features (Figure 8, right graph).

## Discussion

This study aimed at identifying and characterizing an electrophysiological signature of the cortical implication in regulating standing balance. It revealed that during challenging standing conditions, postural sways are represented in ongoing SM1 cortical activity, in the form of CKC at 1–6 Hz. Moreover, delay estimation indicated that cortical activity led CoP excursion by ∼45 ms, while CoP velocity led cortical activity by ∼45 ms, indicating the existence of both afferent and efferent contributions to CKC in the context of balance regulation. Finally, the level of CKC correlated positively with the degree of increase in instability induced by task complexity. Overall, our results indicate that CKC reflects the closed-loop brain–peripheral interaction processes by which SM1 cortical areas intervene to maintain balance.

### Interpretation of the coupling between CoP and cortical activity

Maintaining an upright stance has long been known to be a highly dynamic task. Owing to its upright unstable position, the standing human body is in constant movement, known as postural sways, which are monitored by multiple senses for correction.^6^ Our study clarifies the nature and underlying mechanisms of the cortical contribution to these monitoring and correction processes. That is, we have found that fluctuations of the CoP are represented in sensorimotor cortical activity, at frequencies below 10 Hz, mainly between 1 and 6 Hz. Such cortical representation of movement kinematics, CKC, has been reported in a multitude of motor tasks, ranging from repetitive movements of the upper or lower limb,^21,22,26^ to isometric pinch contraction where a coupling is seen with below 3-Hz minute fluctuations in position.^19^ Our results therefore confirm that CKC reflects a general mechanism for motor control, also involved in maintaining balance.

In our study, CKC was very low or absent during regular standing and became clearly visible in the most challenging condition consisting of standing on foam eyes closed. In unperturbed conditions, balance is thought to be regulated mainly by spinal and brainstem structures,^5^ with minimal implication of the cortex, in agreement with CKC being undetectable in most of the participants (<25 % per feature) in such conditions based on 10 minutes of EEG recordings. When standing on a foam block, the ability to sense pressure distribution and body orientation decreases,^27^ while the effectiveness of ankle torque produced for postural stabilization is reduced,^28,29^ leading to an increase in muscle activity.^30^ Closing the eyes further suppresses visual information, leaving only vestibular information intact to maintain balance. As a result, a greater degree of cortical involvement seems necessary,^5^ and this is precisely reflected in the appearance of significant CKC in our data.

Altogether, the pattern of variation of CKC across conditions strongly indicates that it reflects the degree of cortical involvement in balance regulation.

CKC was higher for CoP features computed along the AP axis compared to the ML axis. This was expected considering the bipedal nature of the task known to mainly recruit lower limb flexor and extensor muscles, especially those acting across the ankle joint.^31^

Previous electrophysiological assessments of balance maintenance have focused on corticomuscular coherence with mixed results on its very existence in that context, and on whether it is modulated by condition complexity.^2,4,32,33^ And although corticomuscular coherence and CKC can be considered as independent and complementary means to probe sensorimotor control,^20^ our data indicates that CKC possesses more desirable features. First, it appears to be relevant for balance maintenance as its amplitude correlated with the relative increase in instability. And second, significant CKC with at least one CoP feature could be uncovered at the individual level, in 97% of our participants in the most challenging condition. Moreover, if CKC indeed scales with perceived task complexity, it is reasonable to think that it could be successfully recorded in an even larger proportion of individuals when balance is further challenged by task complexity (e.g., standing on one foot), pathology (e.g., vertigo), or aging. Therefore, CKC appears to be a highly promising neurophysiological marker of cortical involvement in balance maintenance, for use in future fundamental and clinical research endeavors.

With the advent of EEG technology and signal processing,^34^ a growing amount of EEG studies have moved to studying cortical activity during more natural settings, with a strong focus on gait. These studies have reported significant modulations of the beta rhythm and corticomuscular coherence along gait cycles.^35,36^ And although some also reported significant coupling with muscle activity at lower, <8 Hz frequencies,^20,36^ gait-induced artifacts make it challenging to tell to which extent this coupling arises from genuine brain signals. In that regard, EEG investigation of standing balance through CKC appears to strike the right balance between ecological validity and feasibility. That is, it nicely introduces the complexity of a full weight-bearing task requiring whole body coordination and moving away from well-controlled but too simplistic assessment of uniarticular movements,^37,38^ while at the same time minimizing motion and related artifacts.

### Afferent and efferent contribution to CKC

Our directionality results suggest that both afferent and efferent pathways contribute to CKC, with a ∼45-ms delay from AP CoP velocity to cortical activity and from there, a ∼45-ms delay back to CoP excursion. Importantly, this pattern of results was not related to a mere ∼90-ms delay in between vCoP_AP_ and rCoP since this delay was deemed not significantly different from 0, and since vCoP_AP_ and rCoP showed a very limited degree of coherence in the 1–6 Hz range. Therefore, our results point at the existence of a closed-loop regulation of CoP, where the brain would monitor the CoP velocity and control its position.

The 44-ms delay in monitoring the velocity is roughly compatible with that of 50–100 ms previously reported for CKC in SM1 in response to rhythmic movements of the index finger.^23,26^ In the context of upper limb continuous movement, CKC is thought to reflect cortical processing of proprioceptive afferent signaling.^20,23,24^ Therefore, this delay suggests that similar cortical mechanisms are at play in regulating balance, especially in situations of high postural demands such as standing on foam. Accordingly, the brain would monitor CoP velocity, which should accurately reflect the rate of change in length of the muscle spindles within muscles acting across the ankle joint. However, it should be noted that any derivative of the CoP in the AP direction (position, velocity or acceleration) should lead to identical results, simply because coherence is not affected by a linear transformation, which time derivation is in the Fourier domain.^39^ A predominant monitoring of velocity would however be compatible with previous results on sensory reweighting, where perception thresholds for the visual, vestibular and proprioceptive systems depend on the velocity of sway.^40^ It would also be in line with modelization-based studies who have highlighted the role of velocity information rather than position or acceleration for the control of postural sway.^41,42^ In any case, our results suggest that CKC derived from CoP velocity provides a marker of the integration of proprioceptive information to inform postural control.

The 47-ms delay in controlling CoP excursion is compatible with delays estimated between brain and lower limb muscle activity during isometric contractions.^43^ It is a bit shorter, though not fully incompatible with the report of a motor evoked potential ∼100 ms before a peak in CoP ML displacement during eyes closed stance.^3,13^ From a biomechanics standpoint, the distance to a neutral CoP position, rCoP, directly scales with the motor correction that would be required from the muscles acting on the ankle and hip joints to align the center of mass with the CoP.^44^ Therefore, supported by the predictions of biomechanics, and in line with the study by Varghese et al. (2015), our results indicate that CKC derived from rCoP reflects the efferent commands sent to muscles for postural control.

### Behavioral interpretation

Our results revealed an overall positive relationship between CKC strength in the most challenging condition (standing on foam eyes closed) and the relative increase in instability induced by this challenging condition, compared with unperturbed standing. Interestingly, such a relationship with CKC was not found when considering the uncontrasted instability in the challenging condition. This suggests that CKC does not reflect how much instability there is, but how much instability exceeds that commonly experienced.

Pragmatically, individuals with more unstable posture on foam with eyes closed should be those having more difficulties to leverage vestibular information. Being fully deprived of vision, they could have relied more on proprioceptive information, which was less reliable than usual when standing on foam.^27^ And indeed, standing on an unstable surface such as foam challenges the capacity of the central nervous system to shift the weight of individual sources of afferent information.^45^ Thus, a stronger impact of the hardest condition on balance could indicate inefficient weight shift of the cortex on proprioceptive integration instead of leveraging vestibular information. As a result, in a predictive coding framework,^46^ the discrepancy between perceived and expected proprioceptive input resulted in larger prediction errors for balance, as reflected by high CKC values.

### Limitations and perspectives

We did not attempt to source-localize the cortical generators of CKC in view of disentangling the contribution of somatosensory and motor cortices. Indeed, in EEG, the accuracy of source localization with EEG is at best ∼1 cm for focal cortical sources.^47^ In addition, owing to the ill-posed nature of the inverse solution, source reconstruction cannot resolve or separate the respective contributions of sources in such close-by areas. Future studies using on-scalp MEG could provide further insight into this important question.^48^

CKC was not observed in all participants, and its amplitude was well below the theoretical maximum of 1. However, limited values of CKC were expected, as the cerebral cortex is thought to be involved in postural control only when balance is compromised outside of what is commonly experienced by the individual. Unsurprisingly, in a young and healthy cohort, crossing these margins and getting the cerebral cortex involved in balance control requires a rather challenging task, especially during bipedal stance. This said, in the most challenging condition, CKC reached significance in a rather large proportion of about 4 participants in 5 for 3 CoP features, and in all participants except one with at least one CoP feature. Therefore, our results demonstrate that CKC could be a useful measure to assess the cortical underpinnings of altered balance, with impactful application to assess pathologies such as vertigo and Parkinson’s disease, or aging.

## Conclusion

Our results demonstrate that human sensorimotor cortical areas take part in the control of standing balance in challenging conditions. In doing so, they control the position of CoP, possibly in response to monitored changes in CoP velocity. Importantly, CKC strength could serve as a predictor of postural instability, quantifying the degree of cortical implication in maintaining balance.

## Material and methods

### Participants

Thirty-six young healthy participants (age: 25 ± 3.5 years; weight: 71.7 ± 18.9 kg; 21 women) participated in this study. Volunteers with a history of neurological, muscular, or vestibular disorders were excluded from the study. The study had prior approval by the ethics committee of Erasme university hospital (CCB: B4062021000323, Brussels, Belgium) and all methods were performed in accordance with the relevant guidelines and regulations. The participants gave informed consent before participation.

### Protocol

Participants completed a total of 8 recordings, 2 for each of the 4 conditions. The foam mat used in the foam conditions (Domyos, Decathlon, Villeneuve-d’Ascq, France) measured 39 × 24 × 6 cm and weighed 0.2 kg. The order of the recordings was counterbalanced and each recording lasted 5 minutes. All conditions were performed barefoot, without speaking, arms straight alongside the body, looking straight ahead at a cross on the wall. The position of the feet was standardized to a narrow stance where heels were touching and the line from the heel to the big toe was set at a 20° angle outward for all conditions. Participants were guarded by an experimenter located just behind to prevent falls to the ground. No contact was made between the participants and the experimenter unless a fall would have occurred without assistance. No assistance was required for this cohort.

Participants were monitored for self-reported fatigue and took mandatory breaks of at least 2 minutes between trials.

### Data acquisition

Cortical activity was recorded during the balance tasks with a 63-channel EEG headcap (Waveguard original, ANT Neuro, Hengelo, Netherlands). Electrodes, embedded in a cap, were arranged according to the 10/20 system. Impedances of the electrodes were kept below 50 kΩ using electrolyte gel, and the reference was set at CPz. EEG signals were amplified (Advanced Neuro Technology, Enschede, the Netherlands) and sampled at 1000 Hz. Additionally, ground reaction forces and moments were recorded at 1000 Hz during each condition with a force plate (AccuSway-O, AMTI, Watertown, MA, USA).

### Data pre-processing

EEG and force plate data were imported in Matlab (Mathworks, Natick, MA, USA). After examination of the raw data in all participants, EEG electrodes on the mastoids (M1 and M2) were removed because they featured high amplitude artifacts caused by poor or unstable skin-electrode contact. Thus, signals from the 61 remaining electrodes were kept for further analyses.

EEG data were band-pass filtered between 1 and 90 Hz and notched filtered at 50 Hz using a zero-lag FFT-based filter. Bad channels were identified and interpolated based on the signals of the surrounding electrodes.^49^ EEG signals were then re-referenced to their common average. Time-points at which EEG amplitude exceeded 10 SD above the mean were discarded from further analyses. In addition, 20 independent components were evaluated from the data with Fast ICA to identify and suppress further physiological artifacts.^50^ Independent components corresponding to heartbeat, eye-blink, and eye movement artifacts were visually identified and corresponding signals reconstructed by means of the mixing matrix were removed from the full-rank data.

Force plate data were band-pass filtered between 0.1 and 150 Hz with a fourth order Butterworth filter.

### Data analysis

#### CoP computation

CoP time-series along the AP (CoP_AP_) and ML (CoP_ML_) axes were filtered between 0.1 and 10 Hz. Their single derivative yielded the velocity (vCoP_AP_ and vCoP_ML_). The vector sum at every time step of CoP_AP_ and CoP_ML_ yielded the excursion (CoP) and that of vCoP_AP_ and vCoP_ML_ yielded the velocity (vCoP).

We also estimated the mean speed as the mean of vCoP and the standard deviation of displacement in the AP and ML directions. We will refer to these 3 quantities as the instability parameters.

#### CKC computation

CKC with CoP features (vCoP_AP_, vCoP_ML_, vCoP and rCoP) and coupling between feature pairs was assessed using coherence analysis. Coherence is an extension of Pearson correlation coefficient to the frequency domain, which quantifies the degree of coupling between two signals, i.e. CKC strength, by providing a number between 0 (no linear dependency) and 1 (perfect linear dependency) for each frequency.^39^ In practice, EEG and CoP data were divided into overlapping 2-s epochs with 1.6-s epoch overlap (leading to a frequency resolution of 0.5 Hz). The epochs were then Fourier-transformed to derive a spectrum of coherence between each CoP feature and each EEG signal, following the formulation of Halliday et al.^39^, and using the multitaper approach (3 orthogonal Slepian tapers, yielding a spectral smoothing of 1.5 Hz) to estimate power- and cross-spectral densities.^51^

#### Amplitude of EEG signals underlying CKC

The peak amplitude of the EEG signals underlying CKC was estimated from the coherence and EEG power spectra, for each subject, condition and CoP feature separately. Indeed, coherence, being akin to a squared correlation, indicates the fraction of variance in one signal (here an EEG signal) that is explained by the other signal (here a CoP feature). Therefore, for each frequency bin in the range 1–6 Hz showing significant CKC (see statistical analysis below), we calculated the EEG_SM1_ power spectral density explained by the CoP feature as the total EEG_SM1_ power spectral density multiplied by the coherence level.

This explained EEG_SM1_ power spectral density was converted back to a time-domain amplitude of a sine wave using the square root, after having applied the necessary correction (multiplication by 2 and multiplication by the frequency bin width of 0.5 Hz). Assuming that all EEG sine waves sum in phase, the amplitude of the EEG signal underlying the CKC with the considered CoP feature was estimated as the sum of each individual sine wave amplitude.

#### Time delay estimation

Time delays were estimated from the phase slope of the cross-spectral density between the identified maximal electrode (EEG_SM1_) and CoP feature, using only the data of participants with significant coherence. The phase-slope for a given participant and CoP feature was estimated at frequencies where significant coherence was found within the range 1–6 Hz. A minimum of 6 consecutive significant frequency bins were required to compute a linear regression. A negative-phase slope indicated that the cortical activity was led by the CoP variation.

### Statistical analysis

Statistical analyses were performed using Matlab (Mathworks, Natick, MA, USA).

A threshold for statistical significance of the coherence (*p* < 0.05 corrected for multiple comparisons) was obtained as the 95^th^ percentile of the distribution of the maximum coherence (across 1–6 Hz, and across the SM1 sensor selection) evaluated between EEG and Fourier transform surrogate reference signals (1000 repetitions).^52^ The Fourier transform surrogate of a signal is obtained by computing its Fourier transform, replacing the phase of the Fourier coefficients by random numbers in the range [-π; π], and then computing the inverse Fourier transform.^52^

Normality of the data was assessed using the Kolmogorov-Smirnov test. When the data was found to not be normally distributed, a Box-Cox transformation was applied prior to subsequent analyses.

Possible effects of the balance condition on the number of available epochs used to compute CKC and on CKC at EEG_SM1_ averaged across 1–6 Hz were assessed using one-way analyses of variance (ANOVA). Post-hoc t-tests were performed when significant differences were uncovered.

To assess the behavioral relevance of CKC in the *foam-eyes-closed* condition, we applied a CCA between standardized CoP instability parameters (or their contrast between *foam-eyes-closed* and *solid-eyes-open* conditions) and peak CKC values for rCoP, vCoP_AP_ and vCoP in the range 1–6 Hz. Regularization was used to deal with the limited sample size at our disposition,^53^ with regularization parameters selected through leave-one-out cross-validation. The statistical significance of the final regression model was assessed with permutation statistics, by comparing the correlation value to its permutation distribution (1,000 permutations) obtained after having shuffled instability—but not CKC—values across participants.

## Data availability statement

The data that support the findings of this study are available from the corresponding author upon reasonable request.

## Author Contributions

**Thomas Legrand**: Conceptualization, Methodology, Investigation, Data curation, Formal analysis, Writing – original draft. **Scott Mongold**: Conceptualization, Writing – review & editing. **Laure Muller**: Investigation, Writing – review & editing. **Gilles Naeije**: Conceptualization, Methodology, Writing – review & editing. **Marc Vander Ghinst**: Conceptualization, Investigation, Project administration, Writing – review & editing. **Mathieu Bourguignon**: Conceptualization, Methodology, Formal analysis, Project administration, Funding acquisition, Writing – review & editing.

## Competing Interest Statement

The authors certify having no affiliation, financial agreement or other involvement with any company.

## Notes

### Competing Interest Statement

The authors have declared no competing interest.

### Summary of Updates

As we reconducted some analyses for the present article, we noticed an irregularity in our scheme to correct for the significance of CKC for multiple comparison. That is, Bonferroni correction was applied to the p-values obtained using a maximum statistical approach, which already corrected for multiple comparisons.

